# *Enterococcal* PrgU mitigates PrgB overexpression toxicity by binding to intergenic RNA downstream of the P_Q_ promoter

**DOI:** 10.1101/2020.07.06.190884

**Authors:** Lena Lassinantti, Martha I Camacho, Rebecca J B Erickson, Julia L E Willett, Nicholas R. De Lay, Josy ter Beek, Gary M Dunny, Peter J Christie, Ronnie P-A Berntsson

## Abstract

Efficient horizontal gene transfer of the conjugative plasmid pCF10 from *Enterococcus faecalis* depends on the sex pheromone cCF10, which induces the expression of the Type 4 Secretion System (T4SS) genes controlled by the P_Q_ promoter. The pheromone responsive P_Q_ promoter is strictly regulated to prevent overproduction of the *prgQ* operon, which contains the T4SS, and to limit the cell toxicity caused by overproduction of PrgB, a T4SS adhesin involved in cellular aggregation. PrgU plays an important role in regulating this toxicity by decreasing PrgB production. PrgU has an RNA-binding fold, prompting us to test whether PrgU exerts its regulatory control through binding of *prgQ* transcripts. With a combination of *lacZ* reporter fusion, northern blot, and RNAseq analyses, we provide evidence that PrgU binds a specific RNA sequence within the intergenic region (IGR), ca 400 bp downstream of the P_Q_ promoter. PrgU-IGR binding reduces levels of downstream transcripts, with the strongest decrease seen for *prgB* messages. Consistent with these findings, we determined that pCF10-carrying cells expressing *prgU* decreased transcript levels more rapidly than isogenic cells deleted of *prgU*. Finally, purified PrgU bound RNA *in vitro*, but without sequence specificity, suggesting that PrgU requires a specific RNA structure or one or more host factors to bind its RNA target *in vivo*. Together, our results support a working model where PrgU binding to the IGR serves to recruit RNase(s) for targeted degradation of downstream transcripts.

**Importance:** Bacteria utilize Type 4 Secretion Systems (T4SS) to efficiently transfer DNA from donor to recipient cells, thereby spreading genes encoding for antibiotic resistance as well as various virulence factors. The conjugative plasmid pCF10 from *Enterococcus faecalis*, originally isolated from clinical isolates, serves as a model system for these processes in Gram-positive bacteria. It is very important to strictly regulate the expression of the T4SS proteins for the bacteria, as some of these proteins are highly toxic to the cell. Here, we identify the mechanism by which PrgU performs its delicate fine tuning of the expression levels. As *prgU* genes are present in various conjugative plasmids and transposons, this provides an important new insight into the bacterial repertoire of regulation mechanisms of these clinically important systems.

## Introduction

Hospital acquired (nosocomial) infections and antibiotic resistance are among the largest threats to global health according to the World Health Organization (1). Pathogens often acquire their resistance genes via transferable plasmids and other mobile genetic elements (2). A common opportunistic pathogen in nosocomial infections is the Gram-positive bacterium *Enterococcus faecalis*. Many clinical isolates of *E. faecalis* harbor conjugative plasmids that code for Type 4 Secretion Systems (T4SS) that are responsible for their transmission, as well as other genes encoding for, among other things, resistance to antibiotics and various virulence factors. One of the best-studied *E. faecalis* plasmids is pCF10, a member of the superfamily of pheromone-responsive plasmids. pCF10 codes for resistance to tetracycline from an acquired Tn*925* transposon, and its *prgQ* operon of ~30 kilobases encodes for cell surfaces adhesins including PrgB (also termed Aggregation Substance, or AS) as well as the entire T4SS (Fig. 1A) (3, 4).

**Figure 1.**
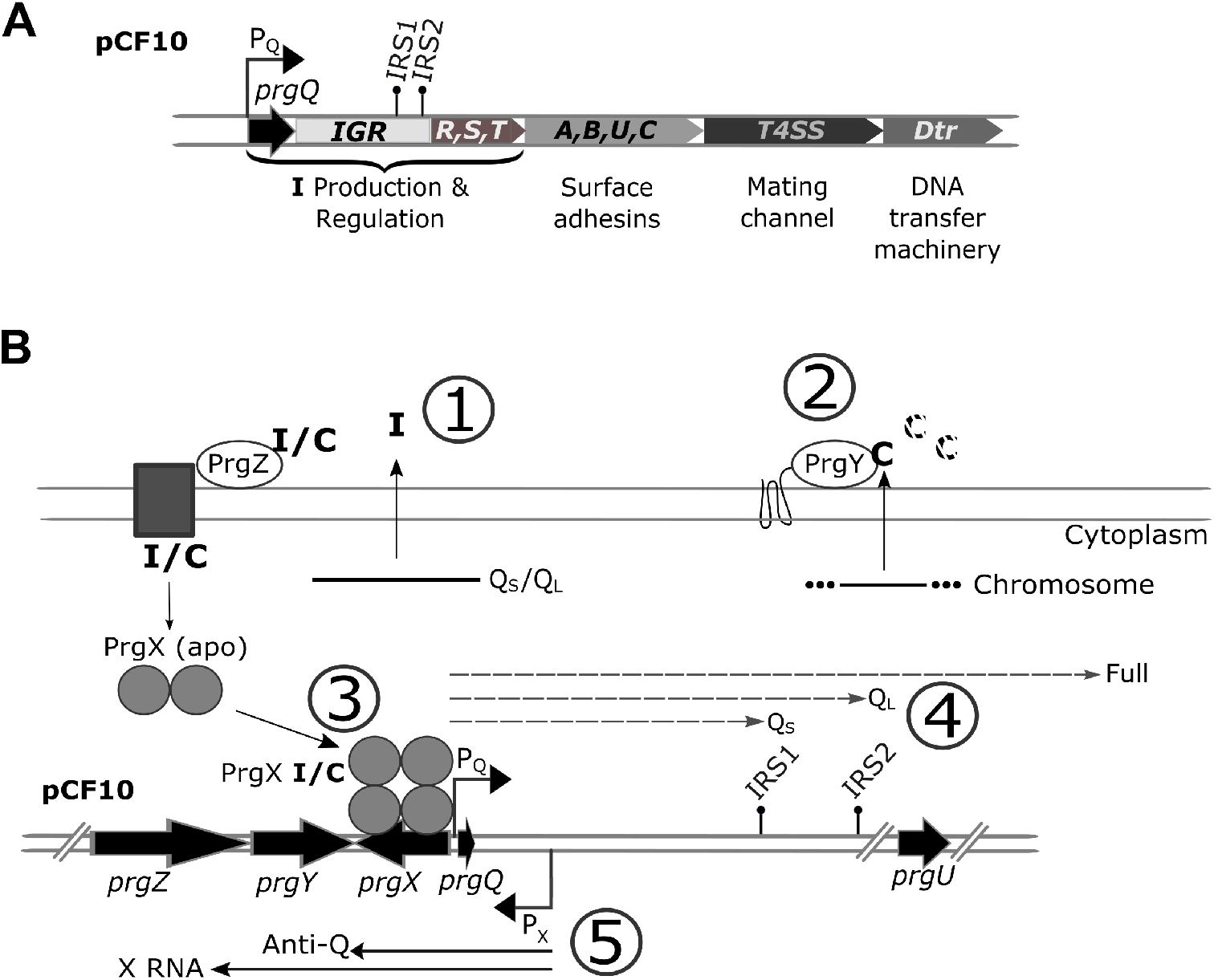
Schematic overview of the *prgQ* operon and its regulation. A) Schematic illustration of the genetic organization of the entire *prgQ* operon (not to scale). B) The inhibitory peptide **I** is transcribed from the pCF10 plasmid (#1), whereas the inducing **C** peptide is transcribed from the genome of all *E. faecalis* cells (#2). Both **I** and **C** are secreted to the outside of the cell, where the **C** peptide from donor cells gets partially degraded by the PrgY protease (#2) (35, 36). Both **I** and **C** are subsequently bound by PrgZ and imported via a permease (20). The transcriptional regulator PrgX is a dimer in its apo state, but tetramerizes upon binding either **I** or **C**. Depending on which pheromone is bound it either represses transcription of the *prgQ* operon (PrgX/**I**) or induces it (PrgX/**C**) (#3) (19, 35, 37). Induction of the *prgQ* operon produces 3 transcripts, Q_S_, Q_L_ and Full (#4) (38). The *prgX* operon also produces two transcripts, one of which is the Anti-Q RNA that aids the formation of a terminator structure of the *prgQ* operon at the inverted repeat sequence 1 (IRS1) (#5) (6). In uninduced cells, Anti-Q levels are sufficient to interact with all nascent *prgQ* transcripts, favoring formation of the IRS1 structure and transcript termination. Pheromone induction leads to the production of many more *prgQ* transcripts, which then overwhelm the pool of Anti-Q. In the unpaired *prgQ* transcripts, IRS1 does not form and transcription extends through the rest of the operon.

The regulation of this system has been the topic of several reviews (3, 4), and an overview is shown in figure 1B. Several negative regulatory checkpoints are involved. Transcription of the *prgQ* operon from the P_Q_ promoter is governed by the ratio of two hydrophobic peptide sex-pheromones; cCF10 (**C** for clumping, sequence: LVTLVFV) and iCF10 (**I** for inhibiting, sequence: AITLIFI). **I** and **C** counteractively modulate the DNA binding activity of PrgX and, in turn, initiation of transcription from P_Q_. The intergenic region (IGR) positioned between *prgQ* and the *prgRST* genes carries two potential transcription terminators, IRS1 and IRS2. Early studies also identified potential secondary structures resembling 23S and 16S rRNA helices in the IGR, which could serve as binding targets for ribosomal proteins (5). Northern blot analysis of pheromone induced donor cells show 2 predominant RNA species, Q_S_ and Q_L_, whose 3’ termini correspond to the position of IRS1 and IRS2, respectively, as well as extended transcripts that include the entire operon (6). A counter-transcript RNA termed Anti-Q regulates formation of the IRS1 terminator structure (6), and it is likely that that IRS1 and IRS2 could also stabilize transcripts produced from 3’→5’ RNase activity, contributing to the total pool of these RNAs. Recently, a small (13 kDa) protein encoded by *prgU* has been shown to play a critical role in controlling over-expression of the *prgQ* operon (7).

PrgU has a pseudouridine synthase and archaeosine transglycosylase (PUA)-like fold as shown by its crystal structure (PDB: 2GMQ). PUA domains are conserved RNA-binding motifs and can be found in all three kingdoms of life (8). They are often found in proteins involved in posttranscriptional modifications of tRNA and rRNA, and in proteins involved in ribosome biogenesis and translation (9). Mutations in PUA motifs are reported to be involved in human diseases such as cancer and dyskeratosis congenita (10, 11). The length of the PUA domains usually varies between 64 and 96 amino acids. In most known PUA containing proteins, these RNA binding motifs are joined to a catalytic domain responsible for RNA modifications (12–14). In contrast, the PUA fold encompasses the entire PrgU sequence (119 amino acids), although PrgU has short insertions in a few loop regions not found in other PUA domains. Deletion of PrgU from pCF10 leads to increased levels of all proteins encoded by the *prgQ* operon (Fig. 1B). This includes PrgB, which is toxic to the cells at high levels (7). Pheromone induction of pCF10Δ*prgU* mutants is therefore lethal if induced for an extended amount of time, and the rare survivors contain suppressor mutations that display a non-inducible phenotype (pCF10∷*prgU*^Res^). It has been suggested that other factors, such as the PrgR/PrgS proteins or bacterial host proteins, could coordinate with PrgU to regulate the expression of the *prgQ* operon (7).

Using a combination of *in vivo* and *in vitro* assays, we show that PrgU regulates expression of *prgQ* operon encoded genes. It does so by interacting with a specific sequence within the IGR, just downstream of the IRS1 sequence. Our cumulative data indicates that this binding depends on another unidentified plasmid- or host-encoded factor(s).

## Results

### PrgU suppresses expression of the prgQ operon

Because PrgU adopts a PUA fold implicated in binding rRNA or tRNA, we hypothesized that PrgU might exert its regulatory control by binding RNA structural folds within the IGR. To test this model, we first constructed reporter plasmids carrying the P_Q_ promoter and a downstream *lacZ* reporter gene. One plasmid (pMC2) contains the IGR in between the promoter and *lacZ*, while another (pMC3) lacks the IGR. A third plasmid (pMC9) contains the IGR (like pMC2), but lacks the *prgX* gene (Fig. 2A). We then monitored *lacZ* expression from these plasmids in induced OG1RF cells (−) or OG1RF cells harbouring pCF10 or pCF10Δ*prgU*, by a β-galactosidase activity assay (Fig. 2B). These strains also carried the vector pDL278p23, which has a constitutive P_23_ promoter (designated P_23_) or a variant of this plasmid, pMB11, that constitutively expresses *prgU* (designated P_23_∷U).

**Figure 2.**
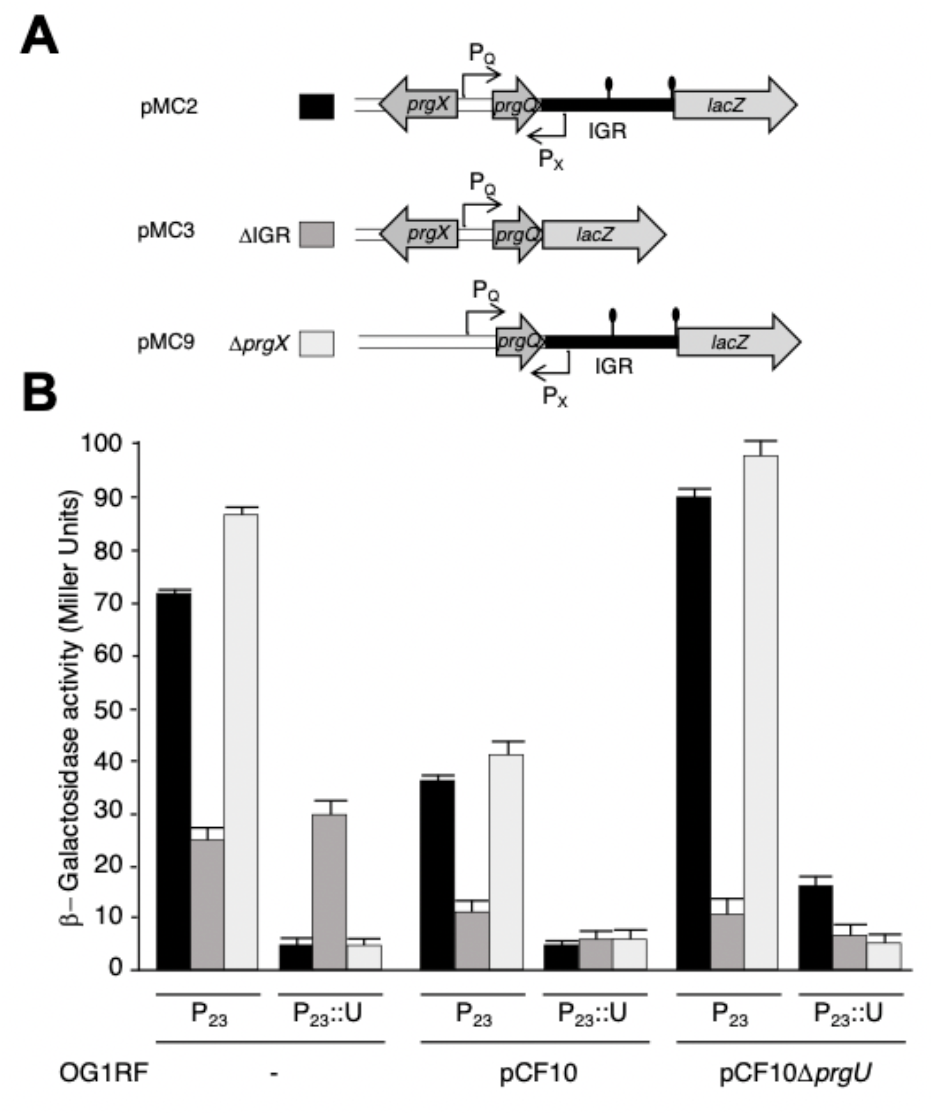
PrgU inhibition of gene expression depends on the intergenic region (IGR) and is independent of PrgX. A) Schematic overview of the plasmids used in this study. The pMC plasmids carry the P_Q_ promoter with its regulatory region and the transcriptional *lacZ* reporter. This reporter gene is positioned at the start-site of *prgR* in pMC2 and pMC9 (Fig. 1B) and at the 3’ end of *prgQ* in pMC3. In pMC9 the *prgX* transcriptional repressor upstream from P_Q_ promoter is deleted. B) β-galactosidase activities originating from expression of the *lacZ* reporter gene on the pMC plasmids listed in panel A (coded in grey scale) in OG1RF cells without a pCF10 plasmid (−), with wild-type pCF10 or pCF10Δ*prgU*. These cells also contain the pDL278p23 vector (P_23_) or this vector constitutively expressing *prgU* (P_23_∷U).

OG1RF cells with the pMC2 plasmid, which contains the IGR (Fig. 2A), and the P_23_ vector plasmid exhibited robust β-galactosidase activity. Isogenic strains that also contains P_23_∷U had only background levels of reporter activity (Fig. 2B, black bars). Introduction of pCF10 into OG1RF(pMC2, P_23_) resulted in lower β-galactosidase activity, presumably due to PrgU production from native *prgU*, whereas reporter levels remained at background levels in cells carrying pCF10 and P_23_∷U due to abundant PrgU production. (Fig. 2B, middle, black bars). As expected, the mitigating effect of pCF10 on β-galactosidase activity was not observed upon introduction of pCF10Δ*prgU* into OG1RF(pMC2, P_23_); in fact, this strain exhibited elevated β-galactosidase activity relative to OG1RF lacking pCF10 altogether (Fig. 2B left & right, black bars).

In cells with the pMC3 plasmid, which lacks the IGR (Fig. 2A), PrgU production from either pCF10 or P_23_:U had no effect on β-galactosidase activities (Fig. 2B compare dark grey bars, P_23_ and P_23_∷U). Cells harbouring pMC3 and pCF10 did exhibit slightly reduced β-galactosidase activities compared to cells with only pMC3. However, this effect was independent of PrgU, as pCF10Δ*prgU* caused a similar reduction of *lacZ* expression from pMC3 (Fig. 2B). We also observed that cells with pMC9, which lacks *prgX*, exhibited slightly higher β-galactosidase activity levels in the absence of PrgU overproduction, as compared to pMC2, which carries *prgX* (compare P_23_ black and light grey bars). This was expected because, in contrast to pMC2-carrying cells, the pMC9-carrying cells completely lack the transcriptional regulator PrgX in the OG1RF background (−) and produce lower levels of the regulator in cells carrying pCF10 or pCF10Δ*prgU*. Importantly, β-galactosidase activity from the pMC9 plasmid was still strongly reduced to near background levels by PrgU overproduction (Fig. 2B, compare light grey bars, P_23_ and P_23_∷U). Taken together, these results establish that PrgU is dependent on the presence of the IGR to inhibit expression of downstream gene(s) and that PrgU acts independently of PrgX.

### Effects of PrgU on Q transcripts in vivo

We next used RNAseq to assess the effects of *prgU* in pheromone-induced donor cells on global, and in particular *prgA*, *prgB* and *prgC*, transcript levels (Fig. 3A, Supplementary dataset 1). The data suggested that the deletion of *prgU* resulted in slightly elevated levels of all transcripts from the *prgQ* operon, but the largest effect was on *prgB* messenger levels (1.9-fold increase in the pCF10Δ*prgU* strain). Although the difference in *prgB* mRNA levels between induced wild-type pCF10 and pCF10Δ*prgU* donors did not reach full statistical significance in this study (Supplementary dataset 1), the effect of PrgU is supported by previous studies where the PrgB protein levels displayed a similar pattern (7).

**Figure 3.**
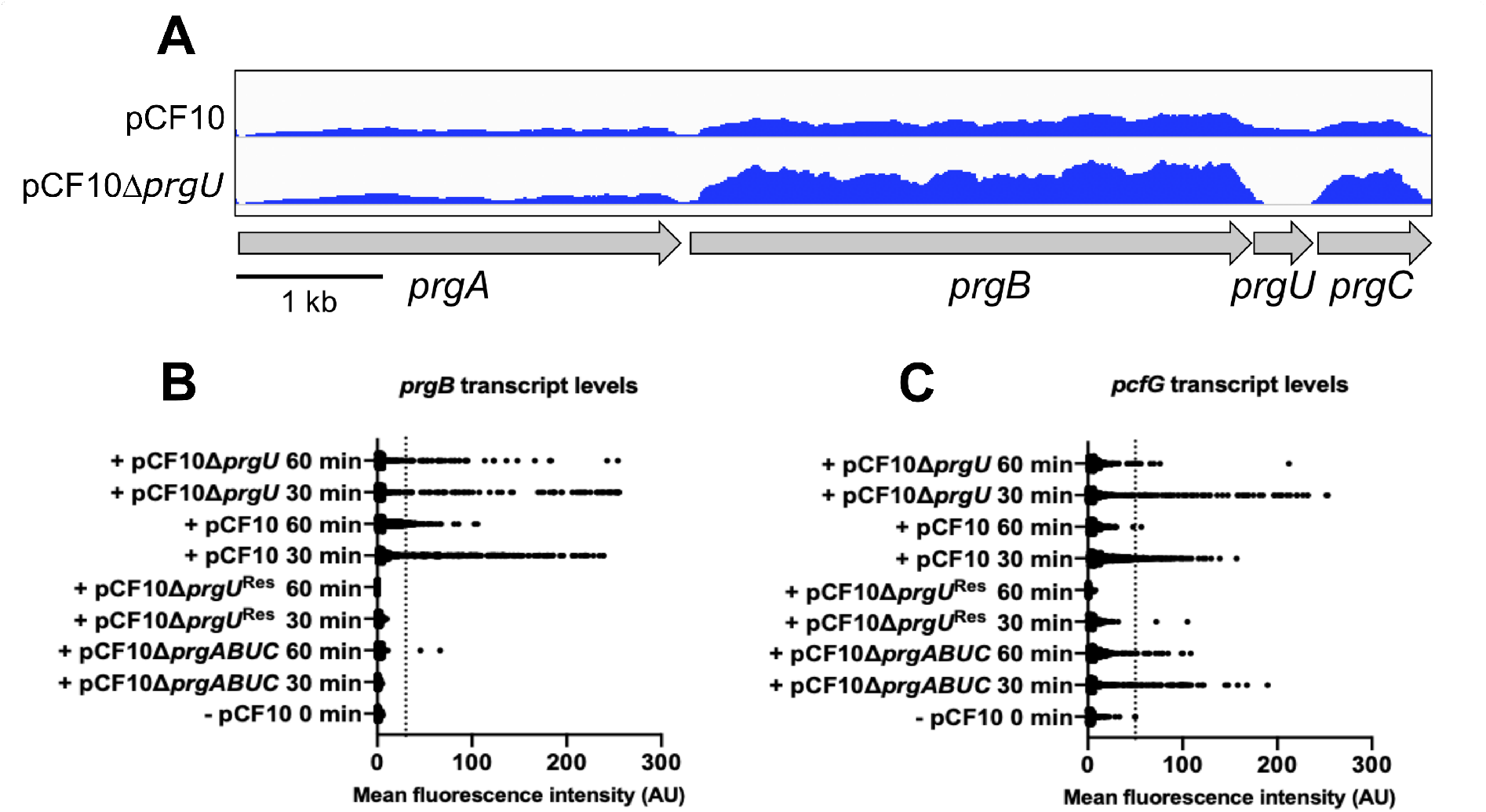
Effects of PrgU on pCF10 transcription at the population- and single cell levels. A) mRNA reads in the *prgA-C* region in induced pCF10 wild-type and pCF10Δ*prgU* cells. The complete RNAseq data for this experiment is available in supplementary data 1. B & C) HCR analysis of *prgB* and *prgG* transcripts in single cells as a quantification of the mean fluorescence intensity per cell of 500 random cells, using hybridization probes against *prgB* (B) and *pcfG* (C) transcripts. Thresholds (thin dotted lines) used are 30 AU and 50 AU respectively. Transcription was induced with (+) 10 ng/mL **C** for 30 or 60 min. Uninduced cells (−) at 0 min and pCF10Δ*prgU*^Res^ containing cells (that cannot be induced) were used as negative controls. pCF10Δ*prgABUC* containing cells are a negative control for *prgB*.

To further evaluate the effect of PrgU on *prgQ* transcript levels, we used single cell *in-situ* hybridization chain reaction (Fig. 3B & 3C). This technique facilitates both quantitative comparison of the levels of different transcripts in the same cell, and also determines the range of variation of expression dynamics within a population that is exposed to the same stimulus (15). The mRNA levels of fluorescently labeled *prgB*, early gene in the operon behind the P_Q_ promoter, and *pcfG,* a late gene in the same operon (11.5 kbp downstream of *prgB*), were examined in single cells. OG1RF cells carrying the following plasmids were analyzed: pCF10, pCF10Δ*prgU*, pCF10Δ*prgU*^Res^ and pCF10Δ*prgABUC*. pCF10Δ*prgU*^Res^ has an additional mutation (*prgR*Δ271) that prevents **C** peptide-dependent induction of P_Q_ transcription (7), and served as a negative control together with uninduced cells (−, pCF10 in Fig. 3B & C). pCF10Δ*prgABUC* was used as an additional background control for *prgB*, as this variant lacks the *prgB* gene, as well as *prgA*, *prgU* and *prgC*.

Cells with a mean fluorescence intensity higher than the threshold for either *prgB* or *pcfG* (Fig. 3B & 3C; thin dotted line, 30 and 50 AU respectively) were considered to be induced. As expected, there was no fluorescence observed for the *prgB* probe (with the exception of two of the 500 randomly chosen pCF10Δ*prgABUC* cells at 60 min), and very little for the *pcfG* probe (only two of the 500 randomly chosen pCF10Δ*prgU*^Res^ cells above the threshold and only at 30 min) in the negative control strains. Also as expected, almost all pCF10Δ*prgABUC* cells exhibited very low fluorescence intensity for *prgB*, but many display high fluorescence intensities for *pcfG*. These controls established that there is no overlap between the two hybridization probes and/or fluorescence channels, consistent with previous findings (16, 17).

To study the effect of PrgU on *prgB* and *pcfG* transcript levels, we compared the levels of fluorescence in cells with pCF10Δ*prgU* to cells with wild-type pCF10. The cells from the pCF10Δ*prgU* strain had a tendency to aggregate and showed an increase in cell lysis, likely due to increased levels of PrgB as previously reported (7, 18). This complicated the measurements in this strain and probably led to an underestimation of the number of induced cells (Table. 1). However, a subset of cells with pCF10Δ*prgU* did display higher levels of fluorescence intensity than wild-type pCF10. This is seen in figure 3B and C by comparing these two strains and is especially pronounced for *pcfG*. Furthermore, the data shows that induced cells with pCF10Δ*prgU* contained a higher amount of transcripts for a longer amount of time than wild type (compare pCF10 and pCF10Δ*prgU* at 60 min). Overall, these findings are consistent with the RNAseq data collected for pCF10Δ*prgU*, shown in figure 3A.

**Table 1.** Percentage of cells that have transcript levels of *prgB* or *pcfG* above the threshold, at 30 and 60 min after induction (from Fig. 3B & 3C). The gray column shows the decrease of cells above threshold at 60 min as compared to 30 min, as a percentage. Note that the number of induced cells is probably underestimated for the pCF10Δ*prgU* strain (see main text).

|  | Percentage induced cells (*prgB*) |  | Decrease (*prgB*) | Percentage induced cells (*pcfG*) |  | Decrease (*pcfG*) |
| --- | --- | --- | --- | --- | --- | --- |
|  | 30 min | 60 min |  | 30 min | 60 min |  |
| pCF10 | 46 | 13 | 73 % | 15 | 0.4 | 97 % |
| pCF10Δ*prgU* | 16 | 9 | 44 % | 14 | 1.6 | 88 % |

Additionally, we observed that all pCF10-carrying cells with high levels of *pcfG* transcripts also had high *prgB* transcript levels, except for the pCF10Δ*prgABUC* containing cells that lack *prgB,* but retain *pcfG*, whereas the converse was not the case (example shown in supplementary figure 1). This was noted previously and could be due to the fact that it takes the RNA polymerase roughly 7 minutes to traverse from the *prgB* gene to the *pcfG* gene (16).

We further investigated the effect of PrgU on *prgQ* operon transcript levels with northern blot assays. As shown in figure 1, transcription from P_Q_ results in the formation of three transcripts, Q_S_ and Q_L_, whose 3’ termini correspond to the IRS1 and IRS2 structures in the IGR, respectively, and the full-length transcript encoding the surface adhesins and T4SS. We sought to compare Q_S_, Q_L_ and full-length transcript levels, but because the full-length transcript from pCF10 could not be resolved due to its length, we instead analysed transcripts generated in strains harbouring pMC2. This enabled detection of the full-length *lacZ* transcript as a proxy for the transcript of the full *prgQ* operon (Fig. 2A). In total RNA extracts of OG1RF(pMC2) cells lacking *prgU*, the Q_S_ and full-length (Full (pMC2)) transcripts were detected, but not Q_L_ (Fig. 4, first lane). However, in extracts from cells expressing PrgU, either from the P_23_ promoter or from the pCF10 plasmid, the Q_L_ transcript was detected. Quantification of the bands showed that the amount of the Q_S_ transcript in P_23_∷*prgU*, pCF10 and pCF10Δ*prgU* increased 3-fold as compared to OG1RF + pMC2. Full-length pMC2 transcripts were no longer detected in the strain that overproduced PrgU from the P_23_ promoter, and the level of this transcript was reduced by ~40% in the strain with wild-type pCF10.

**Figure 4.**
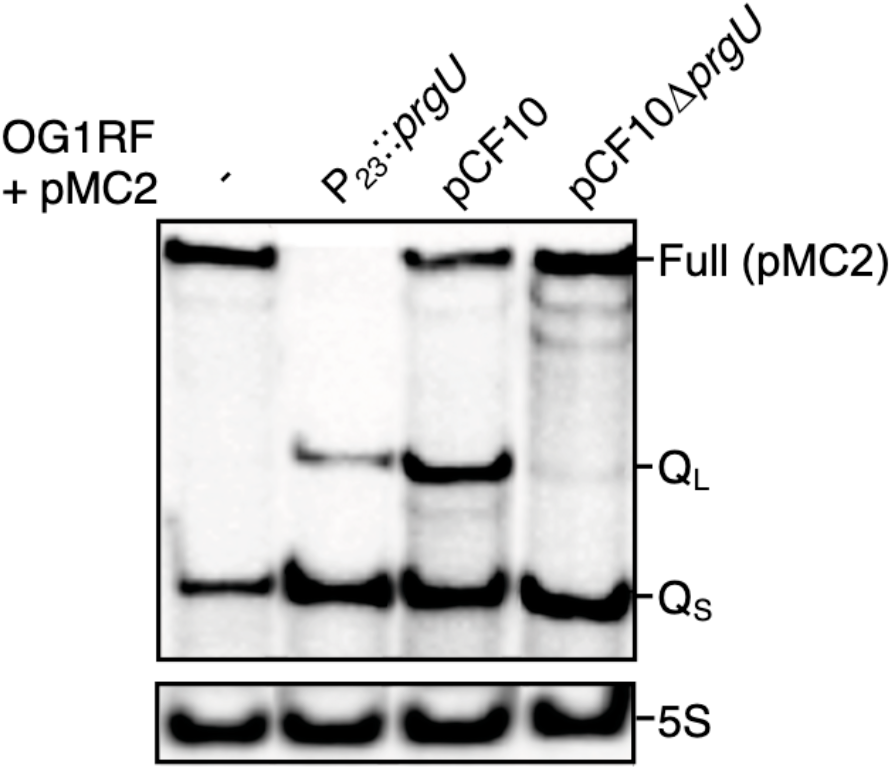
PrgU expression reduces the formation of full-length (Full (pMC2)) transcript and leads to formation of Q_L_ and, at high concentrations, an increase in Q_S_ transcripts. Northern blot analyses of total RNA extracts probed with an oligonucleotide specific for the 5’ end of the IGR, or 5S RNA as a loading control. All strains analysed were OG1RF cells containing the pMC2 plasmid, induced by addition of exogenous **C**. In one strain no additional plasmid was present (−). The other strains either contained the pMB11 vector that constitutively expresses *prgU* (P_23_∷*prgU*), wild-type pCF10 or pCF10Δ*prgU*. Note that Q_S_ and Q_L_ are also formed from the pCF10 plasmids, while the full-length transcript produced from pCF10 is much larger and not visible here.

Taken together, these data suggest that PrgU blocks the formation or enhances the breakdown of RNA transcripts downstream of the IGR. Since PrgU has a PUA-fold that is predicted to bind RNA with rRNA-like structures, we hypothesized that PrgU binds Q_L_, which contains the whole IGR with the IRS1 and IRS2 site and possibly also to Q_S_, which contains over half of the IGR including the IRS1 site. To test this hypothesis *in vivo*, we conducted a pull-down experiment with total cell extracts of OG1RF(pCF10) strains engineered to produce FLAG-tagged PrgU or non-tagged PrgU as a control. RNA recovered from the affinity pull-downs was subjected to northern blot analysis with a probe designed to detect Q_S_ and Q_L_. As seen in Fig. 5A, both Q_S_ and Q_L_ transcripts were identified in the pulled down material in the presence of FLAG-tagged PrgU, but importantly, not with the non-tagged PrgU control. By densitometry, we determined that the amounts of Q_L_ transcripts were the same in the input and affinity-enriched sample, but the level of Q_S_ was reduced 5-fold in the affinity-enriched sample as compared to the input. This suggested that FLAG-PrgU bound and pulled down more Q_L_ (or extended Q transcripts processed back to Q_L_) than Q_S_.

**Figure 5.**
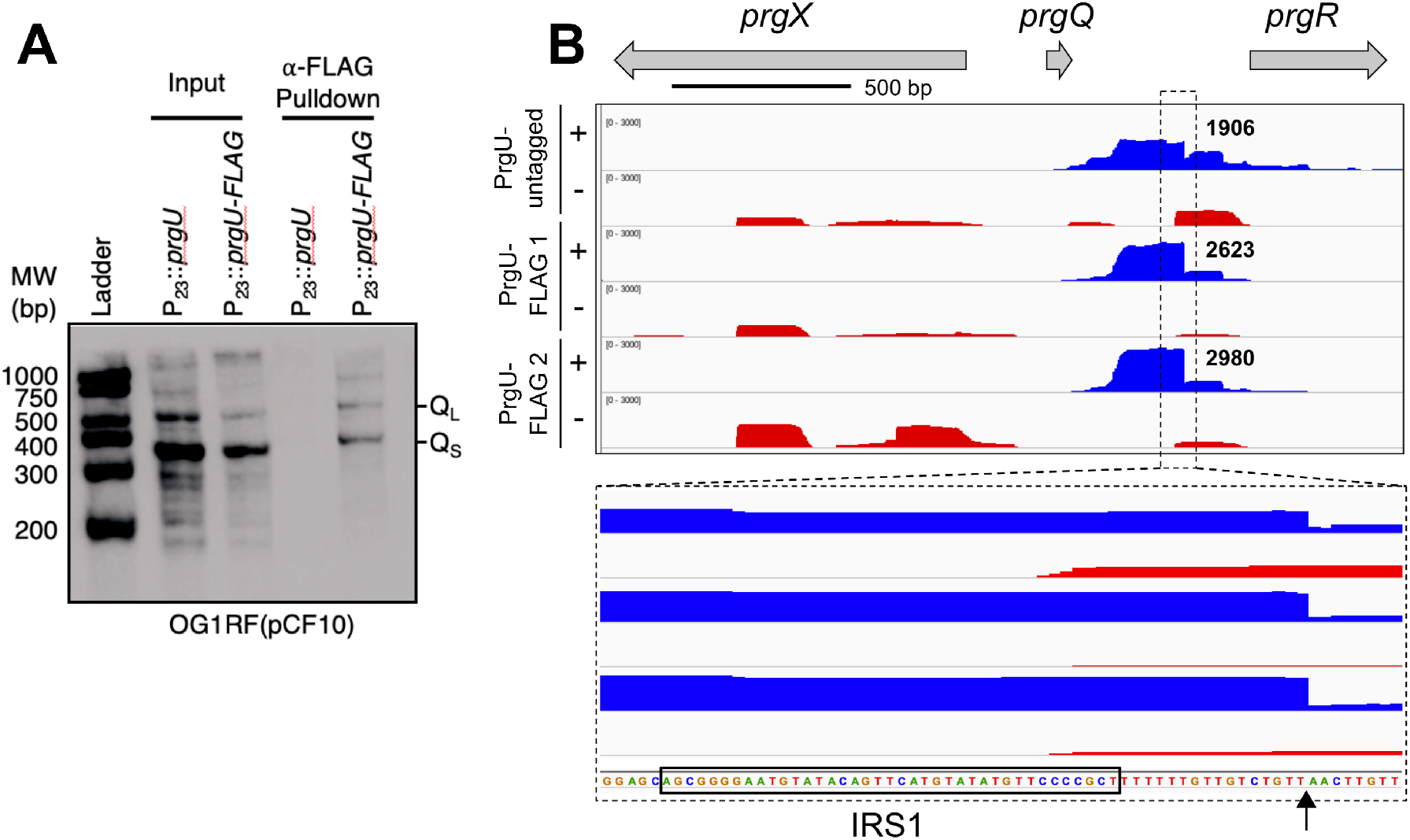
PrgU-FLAG binds to the Q_L_ and part of the Q_S_ transcripts. Cell lysates from **C**-induced OG1RF(pCF10) strains carrying either pMB11 (expresses PrgU from the P_23_ promoter, P_23_∷*prgU*) or pMC10 (expresses FLAG-tagged PrgU from the P_23_ promoter, P_23_∷*prgU-FLAG*) were mixed with magnetic beads coated with FLAG-antibodies. A) Samples from the total cell lysates (input) and the elution fractions from the washed beads (pull-down) were subjected to northern blot analysis using an oligonucleotide protein probe specific for the 5’ end of IGR RNA. Transcripts corresponding to the size of Q_S_ and Q_L_ bound to the magnetic beads in the presence of FLAG-tagged PrgU, but not with the non-tagged control. Note that full-length transcripts of the *prgQ* operon from pCF10 are much larger and not visible here. B) RNAseq analysis of the RNA from PrgU pull-down fractions from lysates of either **C**-induced (+) or non-induced (−) cells. The top panel graphically depicts the relative read counts in the *prgQ* region, whereas the lower panel shows an enlargement with the RNA sequence. The sequence corresponding to IRS1 is highlighted by a black box and the putative 3’ end of the prominent pull-down product is indicated by an arrow.

In separate pull-down experiments, we precipitated the affinity-enriched RNA for analysis by RNAseq. This analysis yielded results similar to those obtained by northern blotting, but provided a more precise analysis of the pulled down RNAs (Fig. 5B, Supplementary dataset 2). Remarkably, a very prominent RNA species with a deduced 3’ end was enriched. This end was located 15 bp downstream from the 3’ end of the IRS1 sequence (note that the IRS1 structure is not generated in non-terminated *prgQ* transcripts).

### In vitro characterization of PrgU

The results of our studies described above, suggested that PrgU binds the Q transcripts. To test this directly, we overproduced and purified PrgU to characterize its biochemical properties in isolation. His-tagged PrgU (15.9 kDa) was produced in *E. coli* and was be purified to homogeneity using a 2-step purification protocol. We then assessed the oligomeric state of purified PrgU-His by GEMMA (Gas-Phase Electrophoretic Molecular Mobility Analysis) and SEC-MALS (Size Exclusion Chromatography-Multi Angle Light Scattering). The GEMMA analysis indicates that PrgU exists both as a dimer and a monomer (Fig. 6A). The SEC-MALS experiment was done with a much higher protein concentration (~1 mg/mL versus 0.01 mg/mL for the GEMMA), and shows almost exclusively a dimer form of PrgU (Fig. 6B). Taken together, these data indicate that PrgU is in a concentration dependent monomer-dimer equilibrium.

**Figure 6.**
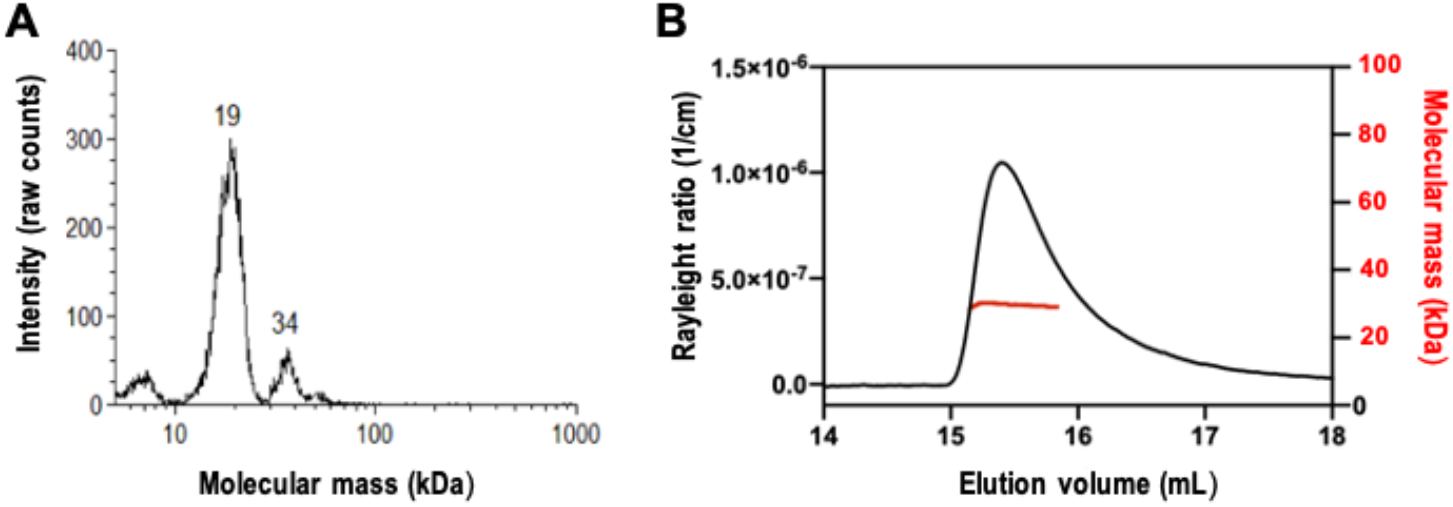
Oligomeric state of PrgU. A) GEMMA analysis of 0.01 mg/mL PrgU. The determined molecular masses are shown above the peaks. B) SEC-MALS profile of PrgU, loaded at a protein concentration of 1 mg/mL, showing a molecular mass of 29 kDa over the range of the peak.

We assayed whether purified PrgU binds double stranded DNA by Electrophoretic Mobility Shift Assays (EMSAs). The results showed that PrgU binds a dsDNA substrate corresponding to the IGR with low micromolar affinity, but that it had a similar affinity for the control dsDNA substrate (Fig. 7A). Next, we investigated the binding of PrgU to ssDNA, dsDNA and RNA forms of the IGR, as well as to shorter sequences corresponding only to the IRS1 and IRS2, by microscale thermophoresis (MST) (Table. S1). However, in the tested experimental conditions, we were unable to detect binding to IRS1 or IRS2 (neither for RNA, ssDNA or dsDNA). We did observe binding to the longer IGR RNA, but PrgU bound equally well to control RNA of similar length (Fig. 7B). Our data therefore suggests that PrgU has low micromolar affinity to both dsDNA and RNA, but that binding is not sequence specific under our experimental conditions.

**Figure 7.**
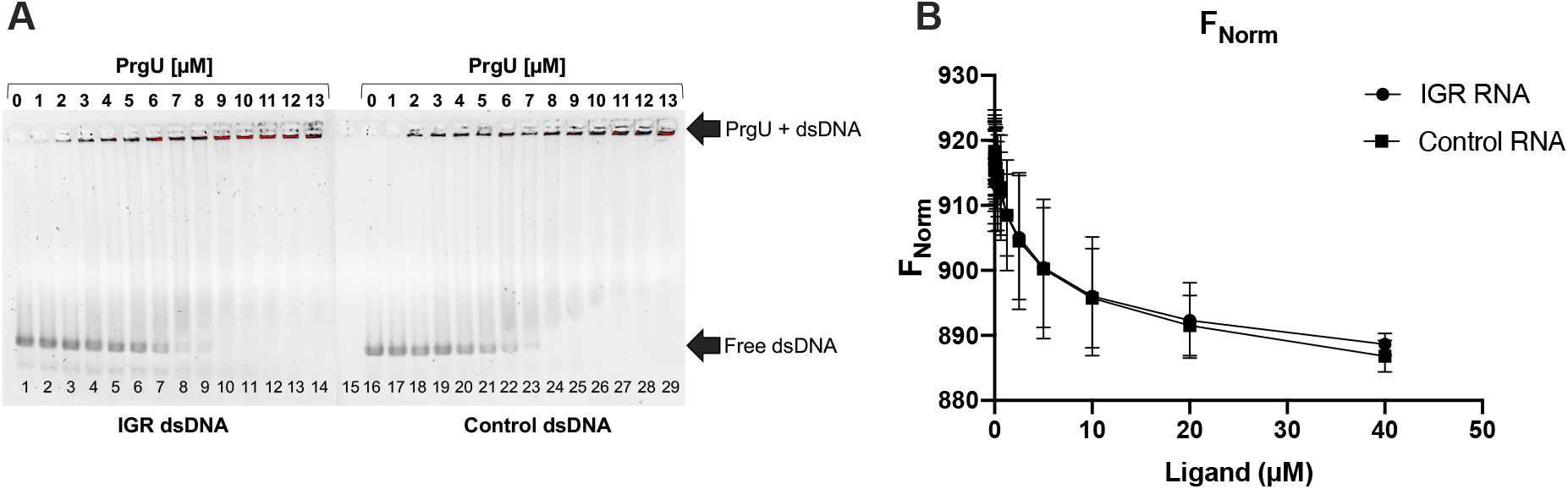
PrgU binding to IGR. A) EMSA with dsDNA of the IGR or an equally long control dsDNA with increasing concentrations of PrgU (0-13 μM). In all cases 100 nM dsDNA was used, IGR (lane 1-14) or control (lane 16-29). Lane 1 and 16 only contains 100 nM DNA, ds-IGR or ds-control, respectively and lane 15 only contains the highest concentration of protein used in this assay (13 μM PrgU). B) Data from MST experiment with a constant concentration of His-labeled PrgU (50 nM), and a varied concentration (between 0-40 μM) of the non-labeled binding partner (RNA IGR and control RNA). n = 3 independent measurements, error bars represent the standard deviation.

## Discussion

Regulation of the conjugative plasmid pCF10 by pheromone sensing has been extensively studied (3, 4, 19, 20). The results have revealed a multitude of mechanisms to regulate the expression from the P_Q_ promoter (3, 4, 19, 21–23). More recently, it was discovered that PrgU is also involved in this regulation. Bhatty et al (7, 18) showed that induction with the **C** peptide was toxic to cells carrying a pCF10 plasmid lacking *prgU* (pCF10Δ*prgU*). This toxicity was relieved by extragenic suppressor mutations or by PrgU overproduction, both of which blocked the production of Prg adhesins. Levels of PrgA, PrgB and PrgC were increased in cells with pCF10Δ*prgU* and, conversely, levels were strongly reduced upon *prgU* overexpression. These findings led to the proposal that PrgU integrates with one or more of the P_Q_ repression systems to control the synthesis of the Prg surface adhesins upon pheromone induction (7).

Here, we showed that PrgU reduces expression from genes downstream of the IGR region (Fig. 2). In our reporter fusion studies and northern blot analyses, we utilized *lacZ* reporter plasmids that lacked all *prg/pcf* genes downstream of IGR. The observed effects of PrgU thus required only the IGR and no other gene products from the *prgQ* operon, including PrgR and PrgS that were previously postulated to coordinate with PrgU to control P_Q_ transcription (Figs. 2, 4) (7). Since none of the reporter plasmids contained a *prgU* gene (Fig. 2), these experiments also confirmed that expression in trans allows for full PrgU function. We also showed that PrgU regulation was unaltered by the presence or absence of *prgX* (Fig. 2), indicating that PrgU does not exert its negative control through effects on formation, stability and/or function of the PrgX/**C** or PrgX/**I** complexes. Based on our findings, we conclude that PrgU functions independently of other known regulators of the pCF10 pheromone response system.

PrgU’s control of expression of the *prgQ* operon depends on the presence of the IGR positioned between the P_Q_ promoter and the regulated genes. PrgU was observed to have a PUA-like fold (7), which is often involved in RNA binding and found in proteins that play a role in posttranscriptional modifications of tRNA and rRNA. This was of special interest, given previous evidence for control of *prgB* expression at both transcriptional and translational levels (5). The data from these previous experiments suggested that ribosomes, sequences in the IGR upstream from IRS1, and a 5’ untranslated region between *prgA* and *prgB* are all involved in a novel translational regulatory mechanism unique to *prgB*. Conceivably this poorly understood mechanism for regulation of PrgB synthesis could be related to the specific effect of PrgU on reduction of *prgB* mRNA and PrgB protein levels.

In addition to data from the transcriptional reporter studies, several other experimental findings support our working model that PrgU exerts its negative control through binding of RNAs produced from the IGR region. We showed that PrgU reduces full-length transcript levels by three independent methods by measuring i) the mRNA levels of *prgB* by RNAseq data (Fig. 3A), ii) the transcript levels from an early gene and a late gene from the *prgQ* operon by HCR analysis (Fig. 3B-C) and iii) northern blot analysis of the RNA from a reporter plasmid (Fig. 4). The northern blot analysis also showed that PrgU overproduction leads to an accumulation of Q_S_ and Q_L_ transcripts, presumably at the expense of the full-length transcript (Fig. 4). Results of the affinity pull-down assays further showed that both of these transcripts (especially Q_L_) co-purified with FLAG-tagged PrgU (Fig. 5A). Furthermore, RNAseq analysis of the affinity-enriched RNA suggests that a predicted linear region just downstream of the IRS1 sequence may be the actual binding target for PrgU (Fig. 5B). We speculate that this RNA was generated by PrgU binding to this region just downstream of IRS1 on much longer transcripts. Bound PrgU would then block 3’ to 5’ degradation by host RNases at this binding site. We also observe the terminated transcript in the non-tagged pull-down (Fig. 5B). It is important to note that PrgU exists also in these cells and thus produces the same terminated transcript as in the cells with FLAG-tagged PrgU. The amount of RNA in the non-tagged sample is less than in the tagged samples, and thus likely represents the identical transcript that bound nonspecifically to the resin during the pull-down experiment.

Both of the reporter constructs (pMC2 and pMC9 shown in Fig. 2) that were negatively regulated by PrgU also contain this predicted PrgU binding site. Interestingly, the essential target region of PrgU is upstream from the *prgU* gene. Such an arrangement could facilitate a timing delay, where PrgU-mediated negative regulation can only take place after a period of protein expression. This would first enable the production of adequate levels of the conjugation machinery for efficient plasmid transfer, followed by a reduction in transcript levels to prevent protein levels rising further and becoming toxic for the donor cells. This is supported by our HCR analysis (Table 1 and Fig. 3B-C).

Despite the several lines of evidence suggesting that PrgU binds a specific RNA target sequence in the IGR, we were unable to confirm that purified PrgU specifically bound the IGR mRNA *in vitro*. EMSA’s with dsDNA of the IGR and control sequences indicated that PrgU non-specifically binds dsDNA with an affinity in the low μM range (Fig. 7A) and RNA with an affinity of ca 2 μM (Fig. 7B). The observed low affinity and sequence nonspecific binding to dsDNA could indicate that PrgU is concentrated around DNA, perhaps associated with active RNA transcription elongation complexes. This would allow it to quickly bind to the specific IGR binding site once it is produced to block further RNA synthesis, or to recruit nucleases involved in RNA turnover. We suspect that specificity for RNA of the IGR could be imparted by another PrgU binding partner, which is missing in our *in vitro* experiments. The candidates include one or more host-encoded regulators or even RNA polymerase itself. Another possibility, raised by Bhatty et al. (7), is that PrgU stabilizes the interaction of Anti-Q RNA or another regulatory RNA with Q-transcripts, which we did not test for in our *in vitro* experiments. We also determined that PrgU exists in a concentration dependent monomer/dimer equilibrium in solution, which might be related to its biological function. PrgU dimerization might enable binding of two RNA target sequences as a prerequisite for effective regulatory control of downstream transcription, or may be necessary for recruitment of co-regulatory protein(s) or antisense RNA.

In summary, we have supplied several lines of evidence supporting our model that PrgU interacts with IGR mRNA and in so doing strongly reduces the production or accumulation of full-length transcripts. Although various mechanisms can be posited to account for PrgU’s inhibition of full-length transcription, two of the more likely ones are: i) transcriptional blockade or ii) recruitment of an RNases(s) for selective degradation. Experiments to determine the exact mechanism of PrgU, as well as the presumptive protein or RNA co-factor, remain exciting avenues for future research.

## Methods

### E. faecalis strains, growth, and pheromone induction

The bacterial strains, plasmids and oligonucleotides used in the study are listed in Table S1 & S2. For plasmid construction, *E. coli* DH5α served as a host. *E. coli* strains were grown in Lysogeny Broth (LB) (Difco Laboratories) at 37°C with shaking. *E. coli* strains were grown with the following antibiotics as needed: chloramphenicol (20 μg mL^−1^), erythromycin (100 μg mL^−1^), spectinomycin (50 μg mL^−1^). *E. faecalis* strains were grown in Brain heart infusion broth (BHI; Difco Laboratories) or M9 minimal media at 37°C without shaking. *E. faecalis* strains were grown with the following antibiotics as needed: erythromycin (100 μg mL^−1^ final concentration for plasmid markers, 10 μg mL^−1^ for chromosomal markers), fusidic acid (25 μg mL^−1^), rifampin (200 μg mL^−1^), spectinomycin (1000 μg mL^−1^ for plasmid markers, 250 μg mL^−1^ for chromosomal markers), streptomycin (1,000 μg mL^−1^), tetracycline (10 μg mL^−1^), chloramphenicol (10 μg mL^−1^). Antibiotics were obtained from Sigma-Aldrich.

### Plasmid constructions

pMC1 is the shuttle vector plasmid pCI372 carrying *prgX* - *prgA* from pCF10 followed by *lacZ* with its own Shine-Dalgarno sequence. It was constructed by PCR amplification of the region of the IGR beginning at a natural *Xba*I site through the end of *prgA* using pCF10 as a template, and amplification of *lacZ* from pCJK205. The two PCR products were digested with *Xba*I-*Bam*HI and *Bam*HI-*Sph*I respectively, and the resulting products were introduced into *Xba*I/*Sph*I-digested plasmid pCIE.

pMC2 is pCI372 carrying *prgX* through the end of the IGR followed by *lacZ*. It was constructed by inverse PCR using pMC1 as a template and the primers listed in Table S2. pMC3 is pCI372 carrying *prgX* through the end of *prgQ* followed by *lacZ*. It was constructed by inverse PCR using pMC1 as a template and the primers listed in Table S2. pMC9 is pMC2 deleted of *prgX*. It was constructed by inverse PCR using pMC2 as a template and the primers listed in Table S2. pMC10 is plasmid pDL278p23 expressing *prgU-FLAG* from the P_23_ promoter; it was constructed by inverse PCR using pMC1 as a template and the primers listed in Table S2.

To produce the His-tagged PrgU overexpression vectors for protein purification, p10-mini (7) was used as a template for PCR amplification. The *prgU* containing PCR products were cloned into the initial cloning vector pINIT using *Sap*I. The constructs were then transferred into p7XC3H via the FX cloning system (24).

### β-galactosidase assays

β-galactosidase activity was assayed as previously described (25) with the following modifications: cells were cultured overnight in M9-YE with selective antibiotics. The overnight culture was diluted 1:5 in fresh medium with 50 ng/mL cCF10 and grown for 90 min to mid-log phase. The cells were pelleted and resuspended in assay buffer (Z buffer: 60 mM Na_2_HPO_4_.7H_2_O, 40 mM NaH_2_PO_4_.H_2_O, 10 mM KCl, 1 mM MgSO_4_, 50 mM β-mercaptoethanol, pH set to 7.0) to eliminate error due to the effects of different carbon sources in the growth medium on the β-galactosidase enzyme activity. Error bars indicate the standard deviation of three replicates in a single experiment.

### Single cell in-situ hybridization chain reaction (HCR)

*E. faecalis* OG1RF cells were used with 4 different plasmids: pCF10, pCF10*ΔprgU*, pCF10Δ*prgU*^Res^ and pCF10Δ*prgABUC* (Table. S2). Overnight cultures were diluted 1:10 in M9 medium and grown to early exponential phase (OD_600_ 1.2). The cultures were induced with 0 or 10 ng/mL cCF10 and harvested after 0, 30 or 60 min.

PFA with a final concentration of 4% was used to fix the cells followed by incubation overnight at 4°C before resuspending in PBS buffer (KPBS) containing RNaseOUT (Invitrogen). HCR labeling was done as described before (17). In short, the cells were permeabilized and HCR labeling was done on cell suspensions. The hybridization probes were designed against an early gene (*prgB)* and a late gene (*pcfG*). The probes and hairpin amplifiers were obtained from molecular instruments (www.molecularinstruments.com). The B2H1, B2H2 and B5H1, B5H2 amplifiers conjugated to alexa flour 647 and 488 were used to detect *prgB* and *pcfG* transcripts. Cells were counterstained with Hoechst 33342.

Cells were then mounted as previously described (17). Images were acquired using a Zeiss Axio Observer.Z1 confocal microscope with an LSM 800-based Airyscan super resolution detector system (Zeiss). Confocal images were acquired through a 63× 1.40- numerical aperture (NA) objective (Zeiss) as z stacks at approximately 0.79 μM intervals using Zen software (version 2.1 Zeiss). Images were deconvolved and flattened using a maximum intensity projection (MIP) with Ortho Display, followed by image analysis by MatLAB (R2019b Mathworks). For HCR analysis the blue Hoechst fluorescence reference channel was used to determine pixel locations of individual cells and co-localized fluorescent overlap from HCR labeled transcripts were quantified. The processed images (deconvolved, flatten by MIP and subjected to background subtraction) were imported into MatLAB (R2019b Mathworks) in a 16 bit TIFF format and analyzed as described previously (17). The reference channel images were binarized using Otsu’s method (26) for thresholding. Objects between 3 and 50 pixels in size were analyzed as cells in the following analysis. HCR intensity value corresponding to each cell was calculated by taking the mean intensity of the pixels corresponding to each cell. To normalize the samples, 500 random cells from each image were used. The induction thresholds were decided based on the expression levels in the uninduced pCF10 population for *pcfG*, while for the threshold determination of *prgB* the pCF10Δ*ABUC* mutant was used. Graphs were created using Graphpad prism 8.0.

### Total RNA extraction from E. faecalis

*E. faecalis* strains were grown overnight in 5 mL M9 media without antibiotics at 37 °C. The culture was then diluted to an OD_600_ of ~0.03 in 50 mL M9 media and grown without shaking to an OD_600_=0.1. Pheromone (cCF10) was added at a final concentration of 20 ng/ml and cultures were grown to an OD_600_=0.3. Cells were pelleted by centrifugation at 16,000 *× g* at 4 °C for 15 min. The pellet was resuspended in 1 mL of RNA-pro solution from a Fast RNA Blue kit (MP Biomedicals) and the mixture was transferred to a fast prep tube containing lysing matrix B. Cells were disrupted by bead beating 5 X at a setting of 400 rpm for 40 sec with 1-2 min intervals on ice. Insoluble material was removed by centrifugation at 16,000 *× g* for 5 min at 4 °C, and the supernatant (~750 μL) was transferred to a fresh microfuge tube and stood for 5 min at room temperature. After chloroform extraction, precipitation with absolute ethanol, and a final wash with cold 75 % ethanol (made with DEPC-treated H_2_O), the RNA pellet was air dried and resuspended in 100 μL of DEPC-H_2_O. The RNA concentration was determined using a nanodrop, and RNA was stored at −80 °C.

### Northern blot analysis

RNA samples (2 μg) were loaded onto a 10% Criterion TBE-urea precast gels (Bio-Rad) and electrophoresed at 70 V until the dye front reached the bottom of the gel. RNA samples were transferred to a Zeta-Probe GT membrane (Bio-Rad) using a Trans-Blot SD semidry transfer apparatus (Bio-Rad) according to the manufacturer’s guidelines. Transferred RNA was UV cross-linked and hybridized overnight with 100 ng/mL of 5’-biotinylated probes (Table S2) in ULTRAhyb hybridization buffer (Ambion) at 42 °C as described previously (27, 28). Blots were developed using the BrightStar BioDetect kit protocol (Ambion) and imaged with a ChemiDoc MP imager (Bio-Rad). Quantifications of the bands were performed by densitometry tracing using Image J software.

### PrgU-FLAG affinity pull-downs

*E. faecalis* strains were grown overnight in BHI (5 mL), and then diluted to an OD_600_ of ~0.03 in 200 mL BHI and grown without shaking to an OD_600_ of 0.1. cCF10 was added at a final concentration of 20 ng/mL and cultures were grown to an OD_600_ of 0.3. Cells were pelleted by centrifugation at 16,000 *× g* at 4 °C for 5 min, washed once with 10 mL TBS buffer (50 mM Tris-HCl, pH 7.4; 150 mM NaCl), and resuspended in 1 mL TBS and transferred to a 1.5 mL eppendorf tube. Cells were pelleted by centrifugation at 16,000 *× g* at 4 °C for 5 min, the supernatant was removed, and the cell pellet was flash frozen in a dry ice-ethanol slurry and stored at −80 °C.

Cell lysates were prepared by suspending cells in 500 μL of TBS buffer with 5 μL of HALT protease inhibitor cocktail and 2 μL of superase. Cells were added to a pre-chilled tube containing 500 μL of glass beads (0.1 mm) and disrupted by repeated cycles of maceration for 40 sec at 400 rpm with intervals on ice. TBS buffer (500 μL) was added and the slurry was vortexed for 30 sec, after which cell fragments and other material were pelleted by centrifugation 12,000 *× g* for 30 min at 4 °C. The supernatant was transferred to a pre-chilled tube, the volume was increased to 1 mL with cold TBS buffer, and 5 μL of superase inhibitor was added.

An aliquot representing the input fraction was removed, and the remaining material was subjected to FLAG pull-downs by addition of 75 μL of anti-FLAG magnetic bead slurry to a 2.0 mL dolphin tube (Costar 3213). The mixture was then placed on a nutator for 2.0 h at 4 °C, after which time the resin was pelleted by centrifugation at 4,000 *× g* for 30 sec at 4 °C. The supernatant was removed, and the resin washed 3 x with 1.5 mL of TBS buffer, with incubations on a nutator for 1 min and then subjected to magnetic bead pull-downs at 4 °C. Material bound to the FLAG beads was eluted by addition of 15 μL 5 μg/μL 3xFLAG peptide solution and 250 μL of 3xFLAG elution buffer. Samples were incubated on a nutator at 4 °C for 30 min, and then centrifuged for 30 sec at 4,000 *× g*. The supernatant containing the eluted material was transferred to a new tube.

TBS (250 μL) and 3 M sodium acetate pH 5.2 (50 mL) were added to the input and the eluted fractions (made as described above), and the RNA was isolated by acid phenol chloroform extraction. The aqueous phase was transferred to a new tube containing 500 μL of neutral phenol/chloroform/IAA, followed by centrifugation at 14,000 rpm for 15 min. The supernatant was then mixed with an equal volume of isopropanol and placed at −80 °C overnight. RNA was pelleted by centrifugation at 14,000 rpm for 15 min, washed once with 75% EtOH, and air dried at room temperature. RNA was suspended in 20 μL DEPC treated water and stored at −80 °C until analysis via northern blot (see above).

### RNAseq analyses

For whole-cell RNAseq, overnight bacterial cultures of OG1RF(pCF10) and OG1RF(pCF10Δ*prgU*) were diluted 1:20 in BHI media and grown for 1 h at 37 °C. Cells were then induced with 10 ng/mL cCF10 followed by incubation for 1 h at 37 °C (29, 30). Ribosomal RNA was depleted using a MICROBExpress kit following manufacturer’s instructions.

For RNAseq on pull-downs, overnight cultures (OG1RF pCF10 carrying either pCIE∷*prgU* or pCIE∷*prgU*-FLAG) were subcultured to OD_600_=0.03 in BHI. At OD_600_=0.1, cCF10 (20 ng/mL) was added. Cultures were grown to OD_600_=0.3 and pelleted (16,000 × g at 4 °C for 10 min). Cells were washed once with 10 mL 1x TBS, once with 1 mL 1x TBS, frozen on dry ice, and stored at −80 °C. Cell pellets were resuspended in 500 μL TBS with 5 μL HALT protease inhibitor cocktail (ThermoFisher) and 2 μL superase, then added to a pre-chilled tube containing 500 μL glass beads (0.1 mm). Tubes were placed in a BeadBug microtube homogenizer (Benchmark Scientific) for 5 cycles (40 seconds, 400 rpm), incubating on ice between cycles. 500 μL TBS was added to each tube, and tubes were pelleted at 12,000× g for 30 min at 4 °C. The supernatant was transferred to a pre-chilled microfuge tube containing 5 μL superase. 50 μL was removed as a protein sample, and 50 μL was removed for an “input” RNA sample. The remaining volume was incubated with 75 μL pre-washed anti-FLAG slurry in a 2.0 mL tube for 2 hr at 4 °C. The resin was pelleted (4,000 × g for 30 sec at 4 °C) and washed with 1.5 mL 1x TBS three times. FLAG-tagged bacterial proteins were eluted with 3xFLAG peptide as follows. 250 μL elution buffer (15 μL of 5 μg/μL 3xFLAG peptide in 500 μL TBS) was added to the resin and incubated with rotation at 4 °C for 30 min. After centrifuging the resin (4,000 × g for 30 sec at 4 °C), the supernatant (containing eluted protein) was transferred to a new tube to which 50 μL 3M sodium acetate pH 5.2 was added. RNA was extracted using with 1 volume acid phenol chloroform. The aqueous phase was transferred to a new tube containing 500 μL neutral phenol/chloroform/isoamyl alcohol (25:24:1), extracted with 1 volume isopropanol, precipitated at −80 °C overnight, and washed with 1 volume 75% ethanol. The pellet was dried and resuspended in 20 μL DEPC-treated water.

All RNA samples were treated with Turbo DNase (ThermoFisher, rigorous method described by the manufacturer) and submitted to the University of Minnesota Genomics Center for Illumina library prep. Whole-cell RNA samples were prepared using a non-stranded TruSeq RNAv2 kit, and libraries were size selected to generate inserts of 200 bp. Pull-down samples were processed using the Illumina TruSeq stranded mRNA preparation kit. To retain small RNAs, size selection was not performed between library preparation and sequencing for pull-down samples. All samples were sequenced as paired-end reads on an Illumina HiSeq 2500 in high output mode (125 bp read length for whole-cell samples and 50 bp read length for pull-down samples). Sequencing files were trimmed to remove contaminating adapter sequences and low-quality bases using Trimmomatic. Reads were mapped to the pCF10 reference sequence (NC_006827) using Rockhopper (31, 32) and visualized using IGV. The RNAseq data can be found in Supplementary datasets 1 & 2.

### Protein production and purification

His-tagged PrgU was produced in *E. coli* BL21(DE3) and grown in terrific broth (TB) media in a LEX bioreactor (Epiphyte). The cells were grown at 37 °C to an OD_600_ of 1.0-1.5, and the temperature was then lowered to 18 °C before induction with 0.5 mM IPTG. The cells were incubated for 18 hrs at 18 °C before harvested via centrifugation and lysed in lysis buffer (50 mM KPi (pH 7.8), 150 mM NaCl, 15 mM Imidazole (pH 7.8) and 10% glycerol) using a Constant Cell disruptor (Constant Systems) with a pressure of 25 kPsi. Cell lysates were run over an immobilized metal-ion affinity chromatography (IMAC) with NiNTA beads using gravity flow and washed with 20 column volumes (CV) of wash buffer (50 mM KPi pH 7.8, 500 mM NaCl, 50 mM Imidazole pH 7.8) followed by 5 CV wash buffer containing 2M Li-Cl followed again by 20 CV wash buffer. The proteins were then eluted with elution buffer (50 mM KPi pH 7.0, 500 mM NaCl and 500 mM Imidazole pH 7.0). Eluted material was run over a size exclusion chromatography (SEC) in 20 mM KPi (pH 7.0), 150 mM NaCl using either a S200 10/300 GL Increase (GE Healthcare) or a S75 10/300 GL (GE Healthcare) column.

### Gas-phase electrophoretic mobility macromolecule analysis (GEMMA)

GEMMA on PrgU was performed as previously described (33). Briefly, the main peak of PrgU after SEC was collected and diluted to a concentration of 0.01 mg/mL in 20 mM ammonium acetate, pH 7.8. The sample was scanned 3 times to increase the signal to noise ratio.

### Size-exclusion chromatography coupled to multi-angle light scattering (SEC-MALS)

200 μL of purified PrgU at 1 mg/mL was run over a Superdex 200 10/300 GL Increase column on an ÄKTA Pure system in line with Wyatt Treos II (light scattering) and Wyatt Optilab T-Rex (refractive index) in 20 mM potassium phosphate (pH 7.0), 150 mM NaCl. Astra software (Wyatt Technology, version 7.2.2) was used to collect and analyze the SEC-MALS data.

### DNA and RNA Oligonucleotides used for in vitro binding studies

The template used for producing the DNA of the IGR and the control was pCF10 or pRS01 respectively (Table. S2). A touch down PCR was done with Phusion polymerase and a starting annealing temperature of 72 °C with a decrease of 1°C /cycle. The PCR products were cloned into pRAV23 (Addgene) using *Eco*RI and *Hind*III restriction sites and transformed into Top10 cells. The cells were grown followed by miniprep (Qiagen) for plasmid isolation. The plasmids were digested with *Eco*RI (Thermo Fisher) and *Hind*III (Thermo Fisher) using the fast digest buffer (Thermo Fisher) and used as a template for T7 *in vitro* transcription. The T7 reaction consisted of 9 mM MgCl_2_, 4 mM rNTPs, 1x Transcription buffer (Thermo Fisher), 2,5-5 μg DNA, 0.05 mg/mL T7 RNA polymerase (Thermo Fisher), and in a 50 μL reaction volume, with incubation at 37 °C for 1 h. The RNA was treated with Proteinase K (Sigma) and DNase I (Sigma) according to manufacturer’s instructions. Afterwards, the RNA was isolated by successive rounds of EtOH precipitation and the dried pellet was resuspended in ddH_2_O at the desired concentrations.

### Electrophoretic mobility shift assay (EMSA)

EMSA was performed as described (34). DNA was from (ThermoFisher) or produced by PCR amplification with pCF10 mini or pRS01 as template (IGR and control DNA). 100 nM DNA was mixed with increasing concentrations of PrgU depending on the experiment. Binding was allowed to occur for 20 min at room temperature (~25 °C) and the complexes were then run at 4 °C on 1% agarose gels at 100 V for 40 min. The visualization was done by post-staining with 3x GelRed and recording on a ChemiDoc (Bio-Rad).

### Microscale Thermophoresis (MST)

The MST was performed as follows. PrgU-His was labeled with the RED 2^nd^ generation His-tag labeling kit (NanoTemper Technologies) according to the manufacturer’s instructions with a final concentration in the assay of 50 nM (molar dye: protein ratio 4:1). A 16:1 dilution series was performed with RNA of IGR and a control sequence of the same length, with 40 μM as the highest concentration in the assay. The samples were mixed in a buffer containing 20 mM KPi (pH 7.0), 150 mM NaCl, 0.05% Tween 20 and 0.1 U/μL Protector RNase inhibitor (Sigma Aldrich) and incubated for 10-20 min at room temperature. After incubation the samples were quickly spun down and loaded into Monolith NT.115 Standard capillaries (NanoTemper Technologies) at room temperature (~25 °C). MST measurements were carried out with a Monolith (Monolith NT.115) using the instrument parameters of 60-100% LED power and medium MST power, the RED filter and an MST on time of 20 s. Data were collected and analyzed using MO.Monolith Control software V1.6.1 and MO.Monolith Affinity Analysis V2.3. Three biological replicates were compared for each sample. Figures were made in Graphpad prism 8.0.

## Acknowledgements

The authors thank Dr. Eric Geertsma for providing the plasmids of the FXCloning system. We would also like to thank Dr. Helmut Hirt for his aid and discussions. This work was supported by grants from the Swedish Research Council (2016-03599), Knut and Alice Wallenberg Foundation and Kempestiftelserna (SMK-1869) to R.P-A.B., and by NIH grants (R35GM118079) to G.M.D. and (R01GM48476 & R35GM131892) to P.J.C.

## Author contributions

L.L. performed protein purification, HCR assays, MST assays and EMSAs. M.I.C. performed *in vivo* analysis of reporter plasmids and northern blots. R.J.B.E. performed HCR assays. J.I.L.W. performed RNA pull-down and RNAseq experiments. L.L., J.t.B., G.D., P.J.C. and R.P-A.B. planned the experiments, performed the data analysis and wrote the manuscript with input from all authors.

## Supplementary datasets, tables and figures

**Supplementary dataset 1:** Differential expression of pCF10 (NC_006827.2) transcripts in OG1RF(pCF10) and OG1RF(pCF10Δ*prgU*).

**Supplementary dataset 2:** pCF10 (NC_006827.2) transcripts identified from PrgU-FLAG pulldowns

**Supplementary table 1:** Oligonucleotides used for PrgU interaction assays.

**Supplementary table 1:** Strains, plasmids, and oligonucleotides used in this study.

**Supplementary Figure 1** Merged microscope image of pCF10 containing cells 30 min after induction showing the fluorescence intensity of *pcfG* transcripts (in green) and *prgB* transcripts (in red) and overlapping cells (in yellow).

